# Comparative effects of central and peripheral PDE10A inhibition on weight gain following semaglutide cessation in diet-induced obese mice

**DOI:** 10.64898/2026.09.07.749896

**Authors:** J.F.E. Grey, B. Whitton, K. Mulvaney

**Affiliations:** BenevolentAI, London, UK

## Abstract

Obesity is a chronic disease of disordered energy balance for which durable weight management remains an unmet need. The rise in use of anorectic therapies, such as incretin analogues, now particularly highlights the need for therapies which maintain weight loss after cessation of effective pharmacotherapy. Phosphodiesterase 10A (PDE10A) inhibition increases energy expenditure and promotes white adipose tissue browning in preclinical models, providing a mechanistic rationale for its therapeutic application. Two complementary studies in male diet-induced obese C57BL/6J mice were conducted to evaluate its pharmacological activity. In a 21-day dose-response study, BEN-8744, an orally bioavailable, peripherally-restricted, PDE10A inhibitor, significantly attenuated body weight gain relative to vehicle at all doses tested. In a 40-day weight maintenance study, mice pre-treated with semaglutide were switched to vehicle-only, BEN-8744, the brain-penetrant PDE10A inhibitor mardepodect or continued semaglutide. Mardepodect was comparable to continued semaglutide, significantly suppressing food intake and weight gain. BEN-8744 showed numerically less gain than vehicle but this difference did not reach statistical significance. Contrary to mardepodect treatment, BEN-8744 also did not significantly improve glucose tolerance or serum insulin. These findings establish BEN-8744 as pharmacologically active through a food-intake-independent mechanism consistent with increased energy expenditure, but indicate that its efficacy is modest relative to additional central PDE10A inhibition. Central PDE10A target engagement is essential for a sustained anti-obesity effect.

## Introduction

Obesity is now recognised as a chronic, relapsing disease of disordered energy balance rather than a consequence of behavioural choice alone. The identification of leptin as a hormone encoded by the ob gene established that body weight is subject to endocrine regulation and that adiposity signals are integrated by hypothalamic circuits to govern appetite and energy expenditure (Zhang et al., 1994). Subsequent work has elaborated the central melanocortin pathway as a principal integrator of these signals, linking peripheral adiposity cues to the coordinated control of food intake and thermogenesis (Yeo et al., 2021). At the tissue level, obesity is characterised by excess and dysfunctional adipose tissue and is driven by a convergence of genetic, neurologic, metabolic, enteric and behavioural processes (Busebee et al., 2023). The global prevalence of adult obesity has risen sharply over recent decades and continues to accelerate across all world regions, imposing a substantial and growing burden of cardiometabolic comorbidity (Phelps et al., 2024).

Despite this burden, durable weight management remains an unmet clinical challenge. Conventional lifestyle and behavioural interventions fail to produce sustained, clinically meaningful weight loss in the majority of patients with obesity (Nguyen et al., 2012). Real-world evidence confirms that most individuals do not achieve recommended weight loss targets even when pharmacotherapy is available. Furthermore, weight regain following cessation of effective therapies, including glucagon-like peptide-1 receptor (GLP1R) agonists such as semaglutide, remains a major limitation (Rubino et al., 2021; Wilding et al., 2022; Budini et al., 2026). There is consequently a pressing need for novel oral agents that can either produce weight loss independently or maintain weight loss achieved by other means, thereby broadening access and improving long-term outcomes.

Phosphodiesterase 10A (PDE10A) has emerged as a promising pharmacological target for chronic weight management. PDE10A hydrolyses both cyclic adenosine monophosphate (cAMP) and cyclic guanosine monophosphate (cGMP), and its pharmacological inhibition by mardepodect (MP-10 / PF-2545920) and THPP-6 have been shown to increase whole-body energy expenditure by hypophagia, and reduce adiposity in preclinical models of diet-induced obesity (Nawrocki et al., 2014; Hankir et al., 2016; Tomaszewski et al., 2023). Mechanistic studies have demonstrated that PDE10A inhibition promotes the browning or beiging of white adipose tissue, enhancing thermogenic capacity in both mouse and human adipocytes (Hankir et al., 2016). These adipose tissue changes have been confirmed in vivo using non-invasive magnetic resonance imaging, further supporting the translational relevance of the pathway (Tomaszewski et al., 2023). Importantly, clinical evidence for the mechanism has begun to emerge. In a randomised proof-of-concept trial of MK-8189, a PDE10A inhibitor developed for schizophrenia, clinically meaningful weight loss relative to placebo was observed as a secondary finding. This provides the first human precedent for PDE10A inhibition as a weight-lowering strategy (Mukai et al., 2024).

## Methods

### 21-day dose-response efficacy study

#### Acclimation and group allocation

Sixty DIO mice were enrolled into a 14-day acclimation period (Day −13 to Day 0). During acclimation, all animals received mock subcutaneous (SC) injections of saline and mock oral (PO) gavage of 1.5% hydroxypropyl methylcellulose (HPMC) vehicle. Twice-daily (BID) dosing acclimation commenced on Day −6 to habituate animals to the BID gavage schedule.

Body weight and food intake were monitored twice weekly throughout the acclimation period. On Day 0, body composition was determined by quantitative magnetic resonance (MRI), a validated technique for non-invasive measurement of fat, lean and fluid mass in living mice (Nixon et al., 2010). Animals were then stratified into five groups of ten (n = 10 per group) on the basis of body weight, food intake and body composition, with ten animals excluded from dosing.

#### Treatment groups and dosing regimen

The five treatment groups were as follows:

1. Vehicle PO BID
2. BEN-8744 0.3mg/kg PO BID
3. BEN-8744 1mg/kg PO BID
4. BEN-8744 3mg/kg PO BID
5. Semaglutide 30nmol/kg SC Q3D

BEN-8744 (BenevolentAI Bio Ltd) was formulated in 1.5% HPMC and administered by oral gavage twice daily for 21 days (Day 1 to Day 21). Semaglutide (Novo Nordisk) was formulated in saline and administered subcutaneously every three days (Q3D) over the same period, consistent with established DIO mouse dosing protocols (Gabery et al., 2020).

#### Endpoints

Body weight and food intake were recorded daily from Day 1 to Day 21. Body composition (fat mass, lean mass and fluid mass) was measured by MRI on Day 0 (baseline) and Day 21 (terminal). On Day 21, animals were killed and terminal blood was collected for serum analysis of total cholesterol (TC), triglycerides (TG), low-density lipoprotein (LDL) and insulin. Adipose tissue depots, namely perirenal adipose tissue (PRAT), epididymal white adipose tissue (eWAT), subcutaneous adipose tissue (SCAT), inguinal white adipose tissue (iWAT) and brown adipose tissue (BAT), were excised and weighed, following established dissection methodology (Bagchi et al., 2019). The primary endpoint was body weight change from baseline.

### 40-day weight maintenance study

#### Acclimation, weight-loss induction phase and re-randomisation

Forty-eight DIO C57BL/6J mice underwent a 2-day acclimation period (Day −2 to Day 0) during which they received mock SC injections of saline at 5 mL/kg once daily. Body weight and food intake were monitored daily. On Day 0, whole blood was collected into EDTA-K2 tubes for baseline glycated haemoglobin (HbA1c) measurement using a Glycated Hemoglobin Analyser. Based on body weight and food intake, 40 animals were selected for dosing.

During the weight-loss induction phase (Day 1 to Day 20), all 40 mice received semaglutide at 30 nmol/kg SC Q3D, together with vehicle (1.5% HPMC) administered PO once daily (QD) on Days 1–7, escalating to BID on Days 8–20. This progressive oral dosing schedule served to acclimate animals to the BID gavage regimen that would be employed in the maintenance phase. Body weight and food intake were recorded daily. Body composition was determined on Day 20 by benchtop NMR (Bruker Minispec LF50).

On Day 20, following 20 days of semaglutide-induced weight loss, the 40 mice were re-randomised into four groups of ten (n = 10 per group) based on body weight, food intake and body composition. This semaglutide lead-in followed by controlled withdrawal design was informed by established paradigms for studying weight regain after GLP-1 receptor agonist cessation in DIO mice (Shah et al., 2025).

#### Treatment groups and dosing regimen (Day 21 to Day 40)

The four treatment groups were as follows:

1. Semaglutide 30nmol/kg SC Q3D + Vehicle PO BID
2. Vehicle PO BID
3. BEN-8744 3mg/kg PO BID
4. Mardepodect 3mg/kg PO BID

#### Endpoints

The primary endpoint during the maintenance phase was body weight change relative to Day 21 assessed daily from Day 21 to Day 40. Body composition (fat mass, lean mass and fluid mass) was measured by Bruker Minispec LF50 benchtop NMR on Day 20 and Day 40 (Nixon et al., 2010). HbA1c was measured from whole blood (EDTA-K2 tubes) at Day 0, Day 21 and Day 40.

On Day 40, an oral glucose tolerance test (OGTT) was performed in all four groups following a 6-hour fast. Mice received an oral glucose load of 2 g/kg at a dosing volume of 5 mL/kg, in accordance with established protocols for high-fat diet-fed mice (Nagy and Einwallner, 2018). Blood glucose was measured at 0, 15, 30, 60, 90 and 120 minutes after glucose administration.

On Day 40, mice were euthanised and blood was collected by cardiac puncture and processed to obtain serum. Serum biochemistry was performed for TC, TG, LDL-cholesterol (LDL-c) and HDL-cholesterol (HDL-c), all measured in mmol/L. Serum insulin concentrations were determined by enzyme-linked immunosorbent assay (ELISA), with insulin reported in ng/mL. Adipose tissue depots (PRAT, eWAT, SCAT, iWAT and BAT) were excised and weighed, with paired depots weighed together; tissues were discarded after weighing.

## Statistical analysis

All data are presented as mean ± SEM. For endpoints measured at a single time point, one-way analysis of variance (ANOVA) was performed followed by Dunnett’s post-hoc test to compare each treatment group against the relevant control group. For endpoints measured over multiple time points (e.g. daily body weight, OGTT blood glucose curves), two-way ANOVA was performed followed by Dunnett’s post-hoc test. Where data did not meet parametric assumptions, the Kruskal-Wallis test was applied. A p value of less than 0.05 was considered statistically significant.

## Results

BEN-8744 is an orally bioavailable PDE10A inhibitor being developed for chronic weight management in adults with obesity. The present study was designed to characterise the preclinical pharmacology of BEN-8744 across two complementary paradigms in DIO mice. First, a 21-day dose-response efficacy study evaluated the dose-effect of BEN-8744 on body weight change from baseline and cardiometabolic indicators. BEN-8744 produced a dose-dependent attenuation of both primary body weight gain and relative weight gain in DIO mice over 21 days, with vehicle-normalised differences ranging from approximately 5.3-9.0% across the 0.3, 1 and 3 mg/kg BID dose groups (Figure 1A-B). Additionally terminal fat mass was also significantly reduced relative to vehicle-only at all doses (Figure 1C). Cumulative food intake across BEN-8744 groups did not significantly diverge from the vehicle-only group (Figure 1D). However on closer inspection of the perirenal, epididymal and inguinal adipose tissues (PRAT, eWAT and iWAT respectively) there was a dose dependent increase which was significant at the highest BEN-8744 dose. In contrast both subcutaneous and brown adipose tissues (SCAT and BAT respectively) did not follow this trend (Figure 1E). Investigation of serum cardiometabolic markers, insulin, triglycerides (TG), total cholesterol (TC) and low density lipoprotein cholesterol (LDL-C) showed that BEN-8744 reduced fed-state insulin and LDL-C at all doses, with the highest dose matching the reduction seen in the semaglutide group (Figure 1F-G).

**Figure 1.**
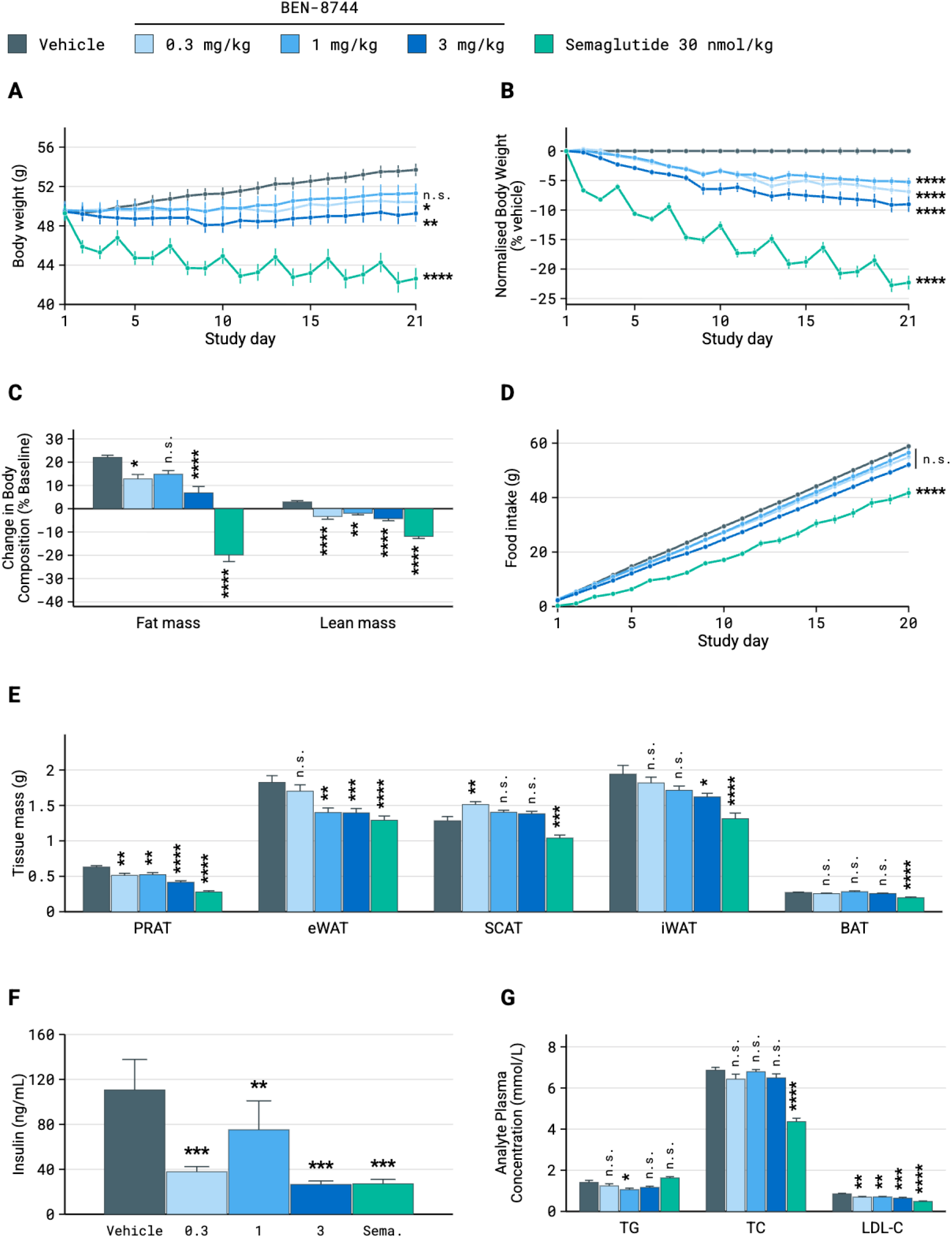
BEN-8744 dose-dependently reduces body weight and adiposity in DIO mice. (**A**) Raw body-weight trajectories. (**B**) Body weight normalised to the contemporaneous vehicle group. (**C**) Change in fat mass (left cluster) and lean mass (right cluster) from Day 0 to day 21, expressed as % of baseline. (**D**) Cumulative food intake. (**E**) Terminal fat-depot mass by tissue (PRAT, eWAT, SCAT, iWAT, BAT). (**F**) Fed-state serum insulin at termination. (**G**) Terminal plasma triglycerides (TG), total cholesterol (TC) and low-density-lipoprotein cholesterol (LDL-C). Statistical comparisons are against vehicle at day 21 (**A, B, C, E**) or at termination (**D, F, G**): *p < 0.05, **p < 0.01, ***p < 0.001, ****p < 0.0001; n.s. = not significant. In (E) the bracket denotes n.s. for all three BEN-8744 dose groups.

In light of blunted weight gain observed in BEN-8744 GLP-1R agonist naive DIO mice, a subsequent 40-day weight maintenance study was undertaken to assess the impact of BEN-8744 in a post-GLP1R agonist paradigm. After a 20-day induction phase with semaglutide, DIO mice treated with BEN-8744 or mardepodect included as a centrally-penetrant comparator were assessed for their impact on weight loss-maintenance and cardiometabolic health. BEN-8744 showed a trend (p_adj_ = 0.13) towards blunted weight gain in the post-semaglutide setting; contrasted by parity between mardepodect and continued semaglutide treatment (Figure 2A). This was matched by vehicle-normalised weight gain, terminal fat mass and lean mass, and cumulative food intake (Figure 2B-D). Furthermore, cardiometabolic readouts were also mirrored between semaglutide and mardepodect (Figure 2E-G‘), whilst BEN-8744 treatment showed a modest but statistically significant reduction in the OGTT at 30 minutes vs vehicle-only (p < 0.01; Figure 2G).

**Figure 2:**
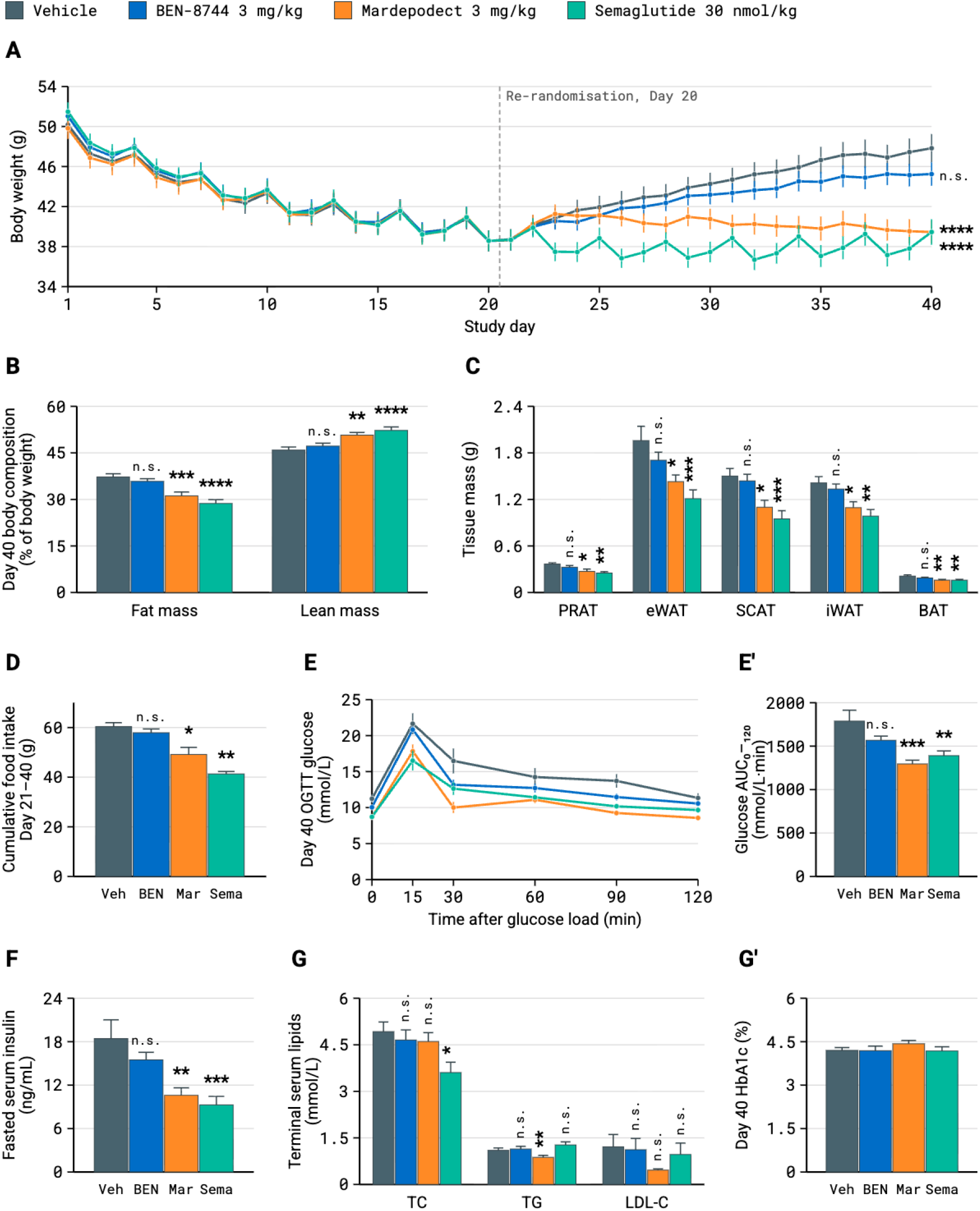
BEN-8744 does not maintain semaglutide-induced weight loss or metabolic improvement after semaglutide withdrawal. (**A**) Body weight, Day 1–40. (**B**) Body composition at Day 40, % of body weight. (**C**) Terminal fat depot mass. (**D**) Cumulative food intake, Day 21–39. (**E**) OGTT at Day 40 and (**E′**) glucose AUC_0_–_120_ □_i_□. (**F**) Fasted serum insulin. (**G**) Terminal serum lipids and (**G′**) HbA1c at Day 40. Significance vs Vehicle: * p<0.05, ** p<0.01, *** p<0.001, **** p<0.0001; n.s. = not significant.

## Discussion

### BEN-8744 efficacy in the 21-day DIO model

The findings here are broadly concordant with the published preclinical literature on PDE10A inhibition in DIO models with a proposed mechanism of action of adipose browning and increased thermogenesis. However the magnitude of effect observed with BEN-8744 was more modest than the pharmacological comparators in these studies, indicating that either the inhibition of the central pool of PDE10A is required for maximal efficacy in the weight-loss setting; or nuances in study design account for discordance in effect drug efficacy (Nawrocki et al., 2014; Hankir et al., 2016; Tomaszewski et al., 2023; Jacobsen et al., 2024). The food-intake-independent weight effect profile is broadly concordant with other non-appetite-based anti-obesity mechanisms, such as TRPM8 agonism (Clemmensen et al., 2018), which produce modest weight loss without detectable changes in food consumption. Additionally, while lean mass declined by approximately 4.2% at the highest BEN-8744 dose, this did not appear disproportionate to published anti-obesity benchmarks which suggest that modest lean mass decline is a typical consequence of pharmacologically driven weight loss (Langer et al., 2026).

### Weight maintenance after semaglutide withdrawal

The weight maintenance study exhibited substantial body weight gain in the vehicle arm consistent with the well-established rebound phenomenon after GLP-1 receptor agonist withdrawal (Rubino et al., 2021; Wilding et al., 2022; Budini et al., 2026). In this context BEN-8744 treatment is consistent with partial weight maintenance, but the study cannot robustly distinguish the BEN-8744 trajectory from vehicle withdrawal. The failure to reach significance may reflect insufficient power, variability in the weight regain kinetics, or a genuinely modest pharmacological effect. This contrasts with other non-GLP1R agonism, peripheral, pharmacological approaches in this setting, namely ATX-304 and ARD-201 (Schneider et al., 2025; Aardvark Therapeutics, 2025). Similarly this contrast was also observed when the central pool of PDE10 was inhibited by mardepodect which was almost indistinguishable in behaviour from continued semaglutide treatment across all readouts, validating previously observed behaviours of CNS-penetrant PDE10A inhibitors in a new context (Nawrocki et al., 2014; Hankir et al., 2016; Tomaszewski et al., 2023).

## Conclusion

This data described here clearly indicates that peripheral PDE10A inhibition alone is insufficient to meaningfully maintain weight loss, insulin sensitivity and improved cardiometabolic health in a post-semaglutide setting. This is likely due to the involvement of the peripheral pool in adipose-derived leptin signalling whereby reduced adiposity, in the post-semaglutide setting, further dampens the effect of peripheral restriction, also explaining blunted weight gain in obese mice only (Tam et al. 2012). However further experiments are needed to confirm this conjecture. Development of PDE10A inhibitors that are centrally penetrant, but avoid neurological adverse effects observed clinically, could offer a novel route for obesity therapy (Megens et al., 2014; Menniti et al., 2021; Mukai et al., 2024).

## Acknowledgements

The work outlined here could not have been done without the work of a vast number of people who worked on, and contributed to, the BEN-8744 programme. Additional thanks to Simona Kolarova for critical review of the manuscript.

## Artificial intelligence Usage Statement

The preparation of this manuscript involved the use of Anthropic’s Claude Opus 4.8, Claude Design and Claude Code. The manuscript was drafted with the BenevolentAI Qor platform.

## References

Bibliography

Aardvark Therapeutics, 2025. Aardvark Therapeutics Announces ARD-201 Preclinical Obesity Data Showing Significant Weight Loss as a Monotherapy, Enhancement of GLP-1RA Therapy in Combination, and Effective Maintenance Following Discontinuation of GLP-1RA Therapy. Press release, 12 August 2025. San Diego, CA: Aardvark Therapeutics, Inc.

Bagchi, D.P. and MacDougald, O.A., 2019. Identification and dissection of diverse mouse adipose depots. Journal of Visualized Experiments, (149), e59499. doi:10.3791/59499

Budini, B., Luo, S., Tam, M., Stead, I., Lee, A., Akrami, A., Vidal-Puig, A. and Park, A., 2026. Trajectory of weight regain after cessation of GLP-1 receptor agonists: a systematic review and nonlinear meta-regression. eClinicalMedicine, 93, 103796. doi:10.1016/j.eclinm.2026.103796

Busebee, B., Ghusn, W., Cifuentes, L. and Acosta, A., 2023. Obesity: a review of pathophysiology and classification. Mayo Clinic Proceedings, 98(12), pp.1842–1857. doi:10.1016/j.mayocp.2023.05.026

Clemmensen, C., Jall, S., Kleinert, M., et al., 2018. Coordinated targeting of cold and nicotinic receptors synergistically improves obesity and type 2 diabetes. Nature Communications, 9, 4304. doi:10.1038/s41467-018-06769-y

Gabery, S., Salinas, C.G., Paulsen, S.J., et al., 2020. Semaglutide lowers body weight in rodents via distributed neural pathways. JCI Insight, 5(6), e133429. doi:10.1172/jci.insight.133429

Hankir, M.K., Kranz, M., Gnad, T., et al., 2016. A novel thermoregulatory role for PDE10A in mouse and human adipocytes. EMBO Molecular Medicine, 8(7), pp.796–812. doi:10.15252/emmm.201506085

Jacobsen, J.M., Petersen, N., Torz, L., et al., 2024. Housing mice near vs. below thermoneutrality affects drug-induced weight loss but does not improve prediction of efficacy in humans. Cell Reports, 43(4).

Langer, H.T., Gilmore, N.K., Hayden, C.M.T., et al., 2026. Weight loss with GLP-1 medicines does not result in a disproportionate loss of muscle mass or function in obese mice and humans. Cell Reports Medicine, 7(3), 102665. doi:10.1016/j.xcrm.2026.102665

Menniti, F.S., Chappie, T.A. and Schmidt, C.J., 2021. PDE10A inhibitors — clinical failure or window into antipsychotic drug action? Frontiers in Neuroscience, 14, 600178. doi:10.3389/fnins.2020.600178

Megens, A.A.H.P., Hendrickx, H.M.R., Mahieu, M.M.A., et al., 2014. Pharmacology of JNJ-42314415, a centrally active phosphodiesterase 10A (PDE10A) inhibitor. Journal of Pharmacology and Experimental Therapeutics, 349(1), pp.138–154.

Mukai, Y., Lupinacci, R., Marder, S., Snow-Adami, L., Voss, T., Smith, S.M. and Egan, M.F., 2024. Effects of PDE10A inhibitor MK-8189 in people with an acute episode of schizophrenia: a randomized proof-of-concept clinical trial. Schizophrenia Research, 270, pp.37–43. doi:10.1016/j.schres.2024.05.019

Nagy, C. and Einwallner, E., 2018. Study of in vivo glucose metabolism in high-fat diet-fed mice using oral glucose tolerance test (OGTT) and insulin tolerance test (ITT). Journal of Visualized Experiments, (131), e56672. doi:10.3791/56672

Nawrocki, A.R., Rodriguez, C.G., Toolan, D.M., et al., 2014. Genetic deletion and pharmacological inhibition of phosphodiesterase 10A protects mice from diet-induced obesity and insulin resistance. Diabetes, 63(1), pp.300–311. doi:10.2337/db13-0247

Nguyen, N., Champion, J.K., Ponce, J., et al., 2012. A review of unmet needs in obesity management. Obesity Surgery, 22(6), pp.956–966. doi:10.1007/s11695-012-0634-z

Nixon, J.P., Zhang, M., Wang, C., et al., 2010. Evaluation of a quantitative magnetic resonance imaging system for whole body composition analysis in rodents. Obesity, 18(8), pp.1652–1659. doi:10.1038/oby.2009.471

Phelps, N.H., Singleton, R.K., Zhou, B., et al. (NCD Risk Factor Collaboration), 2024. Worldwide trends in underweight and obesity from 1990 to 2022: a pooled analysis of 3663 population-representative studies with 222 million children, adolescents, and adults. The Lancet, 403(10431), pp.1027–1050. doi:10.1016/S0140-6736(23)02750-2

Rubino, D., Abrahamsson, N., Davies, M., et al., 2021. Effect of continued weekly subcutaneous semaglutide vs placebo on weight loss maintenance in adults with overweight or obesity: the STEP 4 randomized clinical trial. JAMA, 325(14), pp.1414–1425. doi:10.1001/jama.2021.3224

Schneider, E.J., Hall, J.A., Bor, G., Jacobs, D., Peyer, J.G. and Thieroff-Ekerdt, R., 2025. OR22-05 Weight loss and change in body composition in a DIO mouse model by the combined AMPK and mitochondrial activator, ATX-304, alone, in combination with semaglutide, and after semaglutide withdrawal. Journal of the Endocrine Society, 9(Suppl. 1), bvaf149.080 (ENDO 2025 abstract). doi:10.1210/jendso/bvaf149.080

Shah, H. and Ayala, J.E., 2026. Prolonged semaglutide treatment reveals stage-dependent changes to feeding behavior and metabolic adaptations in male mice. Diabetes, 75(2), pp.288–300. doi:10.2337/db25-0678

Tam, J., Cinar, R., Liu, J., et al., 2012. Peripheral cannabinoid-1 receptor inverse agonism reduces obesity by reversing leptin resistance. Cell Metabolism, 16(2), pp.167–179.

Tomaszewski, M.R., Meng, X., Haley, H.D., Harrell, C.M., McDonald, T.P., Miller, C.O. and Smith, S.M., 2023. Magnetic resonance imaging detects white adipose tissue beiging in mice following PDE10A inhibitor treatment. Journal of Lipid Research, 64(8), 100408. doi:10.1016/j.jlr.2023.100408

Wilding, J.P.H., Batterham, R.L., Davies, M., et al., 2022. Weight regain and cardiometabolic effects after withdrawal of semaglutide: the STEP 1 trial extension. Diabetes, Obesity and Metabolism, 24(8), pp.1553–1564. doi:10.1111/dom.14725

Yeo, G.S.H., Chao, D.H.M., Siegert, A.-M., et al., 2021. The melanocortin pathway and energy homeostasis: from discovery to obesity therapy. Molecular Metabolism, 48, 101206. doi:10.1016/j.molmet.2021.101206

Zhang, Y., Proenca, R., Maffei, M., Barone, M., Leopold, L. and Friedman, J.M., 1994. Positional cloning of the mouse obese gene and its human homologue. Nature, 372(6505), pp.425–432. doi:10.1038/372425a0

